# Stiffness-dependent LOX regulation via HIF-1 drives extracellular matrix modifications in psoriasis

**DOI:** 10.1101/2024.05.14.594058

**Authors:** Parvaneh Balsini, Pauline Weinzettl, David Samardzic, Nina Zila, Maria Buchberger, Philipp Tschandl, Matthias Wielscher, Wolfgang Weninger, Karin Pfisterer

## Abstract

Psoriasis is a common chronic inflammatory skin disease characterized by a thickened epidermis with elongated rete ridges and massive inflammatory immune cell infiltration. It is currently unclear what impact mechanoregulatory aspects in the dermis may have on disease progression. Using multiphoton second harmonic generation microscopy we found that the extracellular matrix (ECM) was profoundly reorganized within the dermis in psoriasis compared to healthy skin. Collagen fibers were highly aligned and assembled into thick, long collagen bundles, whereas the overall fiber density was reduced in psoriasis. This was particularly pronounced within dermal papillae extending into the epidermis. Further, the enzyme LOX, a crucial posttranslational modifier of ECM molecules, was highly upregulated in the dermis of psoriasis patients. *In vitro* functional and knock-down experiments identified a novel link between HIF-1 stabilization and LOX protein regulation in mechanosensitive skin fibroblasts. LOX secretion and activity directly correlated with substrate stiffness, and was independent of hypoxia and IL-17. Finally, scRNA-seq analysis identified skin fibroblasts expressing high amounts of LOX and other ECM-relevant genes and confirmed elevated HIF-1 expression in psoriasis. Our findings suggest a potential yet undescribed mechanical aspect of psoriasis stemming from disordered ECM architecture in the papillary dermis, which could initiate a positive feedback loop in fibroblasts driven by mechanical forces. This mechanism may contribute to tissue stiffening and diminished skin elasticity in psoriasis, potentially exacerbating its pathogenesis.

## Introduction

Psoriasis is an inflammatory skin disease affecting 2-4 % of the global population. The most common form is chronic plaque psoriasis manifesting in reddish, scaly skin lesions that are clearly demarcated from the surrounding normal skin. Psoriasis lesions show epidermal hyperplasia, defective keratinocyte differentiation, increased skin vascularization and infiltration with adaptive and innate immune cells, such as T cells and neutrophils, respectively [1, 2]. The proinflammatory cytokines interleukin (IL)-17, IL-23 and tumor necrosis factor (TNF) have been reported to be the predominant cytokines involved in psoriasis pathogenesis. Biologics drugs targeting these factors for therapeutic interventions have a proven efficiency in up to 80% of affected patients. Why a proportion of patients does not respond to biologics, lose the initial responsiveness over time or show recurring lesions after discontinuing therapy is still elusive [3, 4].

The histopathological features of psoriasis include increased epidermal thickness (acanthosis), increased keratinocyte proliferation, retained nuclei in the corneal layer (parakeratosis), and broad elongated rete ridges that interdigitate with enlarged dermal papillae containing ectatic vessels reaching into the dermo-epidermal junction [3]. Although the stratum spinosum is thickened in psoriasis, the epidermis above the tips of dermal papillae is thinned, increasing skin fragility and resulting in skin fissures when lateral forces are applied. In contrast, both lesional and non-lesional skin is stiffer and less elastic than healthy skin when pulled [3, 5]. Due to the hyperproliferative nature of psoriasis, the massive immune cell infiltration and chronic inflammation, skin homeostasis is often imbalanced because of the high demand of nutrients and oxygen [6]. In particular the main metabolic regulator hypoxia inducing factor 1 (HIF-1) has been shown to be upregulated in psoriasis, but the role of this transcription factor in pathogenesis remains uncertain [6, 7].

Lesions can appear on different body sites and despite ongoing research, it is still unclear what drives their initiation, progression and persistence in some patients, while others are symptom free for years. However, the frequent manifestation of psoriasis plaques at sites with stronger mechanical tensile stress, such as the extensor sites of elbows and knees or the scalp suggests that mechanical signals, force transduction, mechanosensing, conversion into biochemical signals and induction of cell responses may play a role in disease pathology [2, 8]. In addition, the Koebner phenomenon, which is the development of secondary psoriatic lesions upon physical trauma or mechanical stress, further supports the idea that mechanical stress may act as a pathogenic factor in psoriasis [8, 9].

The extracellular matrix (ECM) is a fine-tuned scaffold that enables reciprocal force transduction between tissue cells and matrix components. Fibrillar collagen and elastin fibers are the major structural components within the dermis and responsible for tensile strength and elasticity, respectively [10]. The ECM composition and architecture is regulated quantitatively on a gene expression level and qualitatively via posttranslational modifications such as crosslinking of adjacent ECM fibers. Both parameters define the physical properties of the ECM such as density, pore size, stiffness, elasticity, and permeability for soluble factors regulating cell migration, as well as cytokine and growth factor diffusion and distribution [10, 11]. In particular, collagen fiber topology, orientation and alignment can affect its mechanical properties and thus determine tissue deformation, strength and stability [12]. Tissue cells are able to sense physical parameters such as rigidity, elasticity and stiffness of fibers and adapt to changing environments with cytoskeletal modifications and transcriptional responses, but little is known about dermal cells in this regard.

Fibroblasts are the main producers of tissue-specific ECM components and ECM modifying enzymes such as the copper-dependent enzyme lysyl oxidase (LOX) in the skin [13]. LOX facilitates covalent crosslinks between adjacent collagen fibers resulting in longer fibrils and thicker collagen bundles, thereby changing their mechanical properties [11, 12, 14]. In cancer cells, LOX has been shown to be under the transcriptional control of HIF-1, and has been found to play a role in skin aging, wound healing and fibrosis, but little is known about its role in skin inflammation [15, 16]. LOX is synthesized as a catalytically inactive 50 kDa proenzyme, secreted and proteolytically processed in the extracellular space into the biologically active enzyme of 30 kDa [17].

To date, the impact of mechanoregulatory factors in the skin on disease propagation remains unclear. Here we deep-dived into structural modifications and regulatory mechanisms of dermal ECM with a focus on the chronic inflammatory skin disease psoriasis.

## Results

### ECM modifications in psoriasis

In an attempt to reveal novel mechanisms that drive disease progression in psoriasis, we analyzed scRNA-seq data from healthy and psoriasis skin samples using a recently published dataset from Gao *et al*. (Suppl. Fig. 1) [18]. Differential gene expression analysis using all skin cells, including fibroblasts, pericytes, keratinocytes, melanocytes (MEL), lymphatic endothelial cells (LEC), endothelial cells (EC) and immune cells, revealed that the three top ranked enriched Reactome 2022 pathways were ‘Extracellular Matrix Organization’, ‘Collagen Biosynthesis and Modifying Enzymes’ and ‘Collagen Formation’ (Suppl. Fig. 1A and B). Further analysis of ‘Extracellular Matrix Organization’-associated genes identified several ECM-relevant and -modifying genes dysregulated in cells derived from lesional psoriasis skin when compared to cells from healthy skin (Suppl. Fig. 1C). To validate these findings we next compared bulk gene expression microarray data from Yao *et al*. comprising healthy (H), lesional (L) and non-lesional (NL) skin. We again found many ECM organization-associated genes to be differentially expressed in psoriatic skin when compared to healthy skin (selected genes are shown in Suppl. Fig. 1D) [19]. Some ECM genes were specifically up- or downregulated in L compared to H (COMP and LTBP4, respectively), whereas others were downregulated for both L and NL (ADAMTS9). Surprisingly, we discovered that many ECM-relevant genes were differentially regulated in NL skin compared to H skin, such as genes encoding ECM-modifying molecules, cell-matrix adhesion proteins and collagens that stabilize the epidermal-dermal junction, suggesting an early deregulation of the ECM in psoriasis pathogenesis.

This prompted us to investigate whether the architecture of the ECM is modified in psoriatic skin. Label-free imaging of collagen fibers in tissue sections using Second Harmonic Generation (SHG) microscopy revealed that the ECM was altered in lesional psoriatic skin compared to healthy skin (Fig. 1A). To quantify these changes in ECM organization, we analyzed the overall fiber orientation relative to the skin surface using CurveAlign [20] (Fig. 1B). Collagen fibers were oriented parallel to the skin surface in healthy skin (0° and 180° angle of fibers) compared to a perpendicular orientation in psoriasis (90° angle of fibers, Fig. 1C, Suppl. Fig. 2A). To analyze multiple samples we calculated fiber orientation coefficients, in which perfectly perpendicular fibers (90°) were assigned a value of 1 and fibers that were oriented parallel to the skin surface 2 (0°/180°, Fig. 1D). Healthy skin samples showed a consistent parallel orientation of collagen fibers, whereas collagen fibers in psoriatic skin showed a significant drop in the fiber orientation coefficient, suggesting a high number of fibers oriented perpendicular to the skin surface (Fig. 1D, left plot). Further, the overall fiber density, described as relative fiber density around detected fibers or fiber bundles, was decreased in psoriasis (Fig. 1D, right plot). Finally, we characterized and quantified the fiber parameters length and width by segmenting and identifying individual collagen fibers using the Ridge Detection tool (Fig. 1E) and found that ECM fibers in the dermis of psoriatic skin were significantly longer compared to healthy skin, whereas the overall width was unaffected (Fig. 1F).

**Figure 1.**
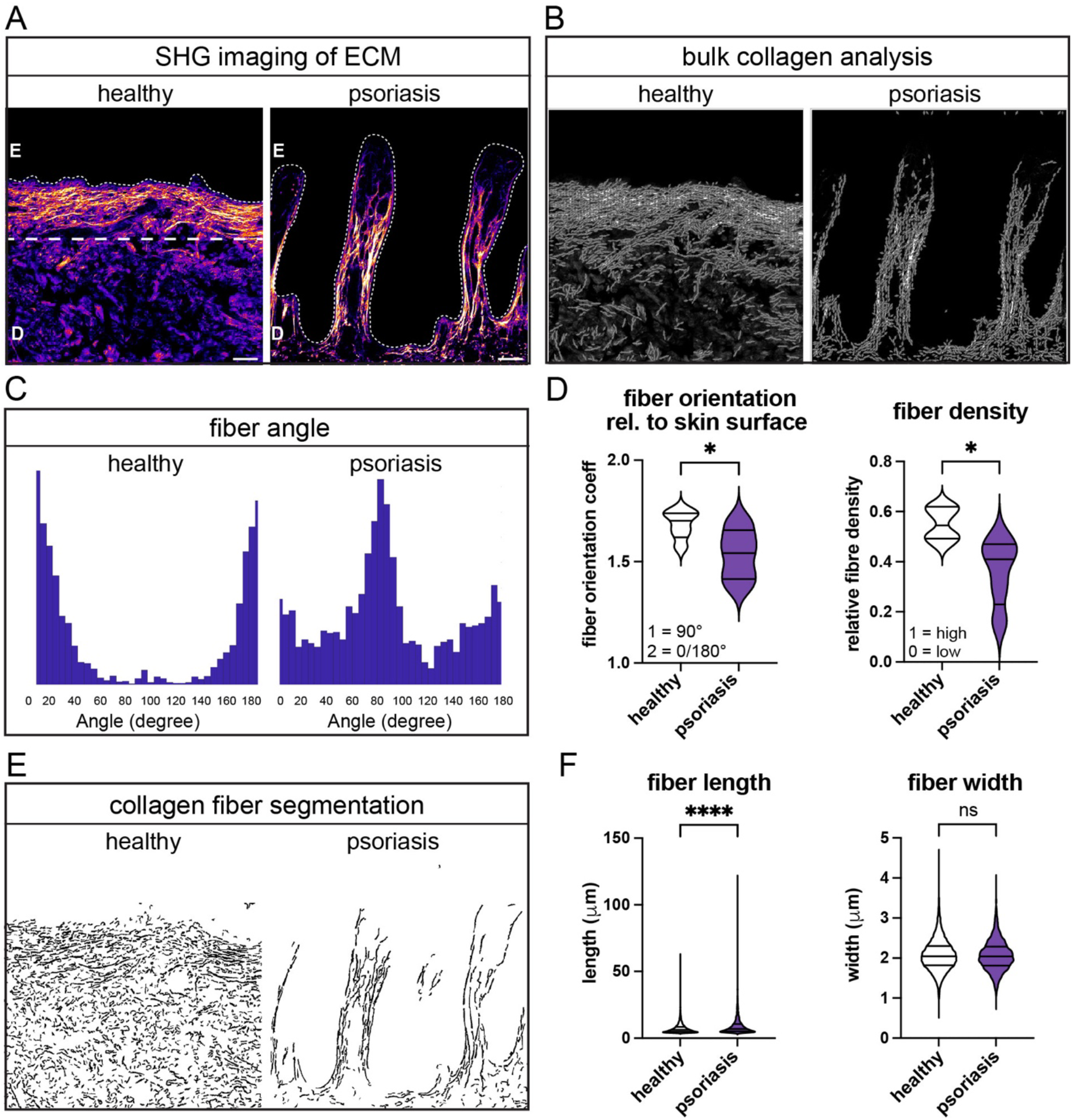
ECM fiber orientation and density are disorganized in psoriasis. **A)** SHG images of the ECM in the dermis of healthy and psoriasis skin. The fine dashed line indicates the epidermal-dermal border. Note that the epidermis is not visible. The thick dashed line indicates the approximate border between papillary and reticular dermis. Scale bar = 50 μm; D = dermis, E = epidermis. **B)** Collagen fibers were detected and used for bulk analysis. Shown are representative segmented fibers from SHG images in A. **C)** Fiber angle histograms for the examples shown in A. **D)** Left plot: quantification of fiber angles for multiple healthy (n=6) and psoriasis skin donors (n=5). Fiber orientation coefficients are shown in which 90° angles were assigned the value 1, and 0 and 180° angles were assigned the value 2. Right plot: shown is the relative fiber density, 1 = high density, 0 = low density. **E)** Shown is a representative segmentation mask for SHG images shown in A) and analyzed with Ridge Detection. **F)** Fiber length and width values were extracted and compared for healthy (n=6) and psoriasis skin (n=5).

### Fiber modifications in dermal papilla

We hypothesized that the ECM is affected differently depending on the dermal compartment. To get an overview of total skin structure in healthy and psoriatic skin, we used Masson-Trichrome staining to detect epidermal keratinocytes (keratin and muscles in red), dermal collagen fibers (blue), cells (cytoplasm in pink) and nuclei (brown) (Fig. 2A). We found a distinct reorganization of the skin’s architecture in psoriasis, on the one hand confirming thickening of the epidermis, enlargement of dermal papillae and increased vascularization, and on the other hand revealing altered collagen-fibril structures, which were particularly pronounced within dermal papillae (insets, Fig. 2A). Collagen fibers built a dense network of fibers in papillae of healthy skin, whereas they appeared less dense and highly aligned in psoriasis. Thus, we analyzed the papillary dermis more thoroughly, in particular dermal papillae and the upper dermis excluding papillae (Fig. 2B). Equal sized regions of interest (ROIs) where selected within both skin compartments of SHG images (Fig. 2C) and fibers were detected using CT-FIRE and single fiber metrics quantified (Suppl. Fig. 2B). Collagen fibers were highly aligned within dermal papillae in psoriasis compared to healthy skin, whereas fiber alignment was reduced in the upper dermis excluding papillae (Fig. 2D). Further within papillae, collagen fibers were significantly longer and wider in psoriasis compared to healthy skin suggesting posttranslational modifications (Fig. 2E).

**Figure 2.**
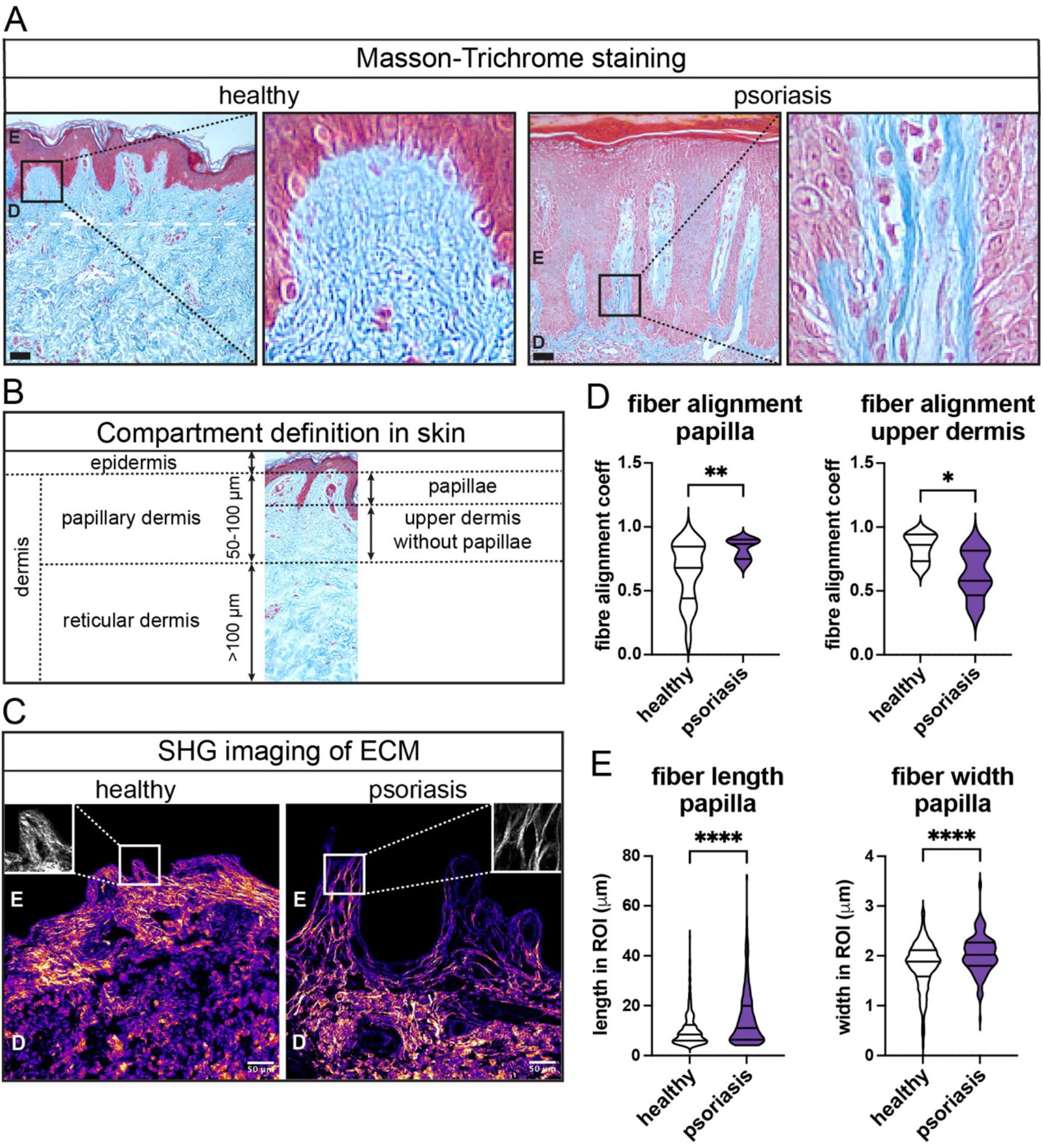
Collagen fiber modifications are most pronounced within dermal papillae. **A)** Masson-Trichrome staining of healthy and psoriatic skin. Collagen fibers are stained blue, whereas cells are purple/pink. Shown is one representative image for each and insets are magnifications of selected dermal papillae. Scale bar = 100 μm; D = dermis, E = epidermis. **B)** Schematics of the organization of human skin into epidermis and dermis, further stratified into dermal papillae and upper dermis within the papillary dermis and the deeper reticular dermis that lays above the hypodermis (not shown). **C)** SHG images of healthy and psoriasis skin with highlighted crop image examples (ROI) that were used for analysis of fiber morphology in papillae. **D)** Fiber alignment values are shown for three ROIs within papillae (left plot, high coefficient = high alignment) and one ROI in the upper dermis without papillae (right plot) for heathy (n=6) and psoriasis skin (n=5). **E)** Fiber length (left plot) and width (right plot) were calculated in ROI crops of papillae in healthy (n=6) and psoriasis skin samples (n=5).

Previous reports suggested that immune cells emerge from dilated blood vessels within dermal papillae and are released by so-called “squirting papilla” [3]. Reoriented ECM fibers may enhance this effect and promote immune cell migration towards the epidermis upon extravasation from the endothelial lumen. Indeed it has been shown that immune cells use ECM fibers for tissue migration [21]. To investigate whether immune cells interact with dermal collagen fibers, we stained CD45+ cells in healthy and psoriasis skin and captured both the ECM and CD45+ immune cells using a multiphoton microscope. Immune cells seemed to colocalize with the ECM in the dermis in both healthy and psoriatic skin (Suppl. Fig. 2C.). However, we observed that immune cells in psoriasis samples were often clustered together encapsulated by ECM (asterisks) and many CD45+ cells attached to elongated collagen fibers within dermal papillae and at the bottom of rete ridges (arrowheads). Hence, reoriented fibers in psoriasis could indeed enable immune cells to relocate to skin compartments they are usually excluded from, such as the epidermis.

### LOX expression is upregulated in psoriasis

Both collagen fiber elongation and bundling of aligned fibers can be induced by posttranslational modifications through matrix modifying enzymes. Lysyl oxidase (LOX) is a key enzyme involved in collagen crosslinking and is implicated in various diseases that entail alterations to the ECM. Immunofluorescent staining for LOX revealed higher expression in psoriatic skin compared to healthy skin (Fig. 3A, left panel). Quantification of LOX intensity in whole skin of multiple donors showed a significant increase of LOX expression in psoriasis compared to healthy skin (Fig. 3B, left plot). *In silico* separation of whole skin into an epidermal and a dermal compartment revealed particularly high mean LOX intensities in the dermis of psoriasis samples, whereas in the epidermis the mean LOX intensity was not significantly increased (Fig 4B). This suggests that LOX may represent a regulator of dermal interstitial matrix modifications and a potential role in psoriasis.

**Figure 3.**
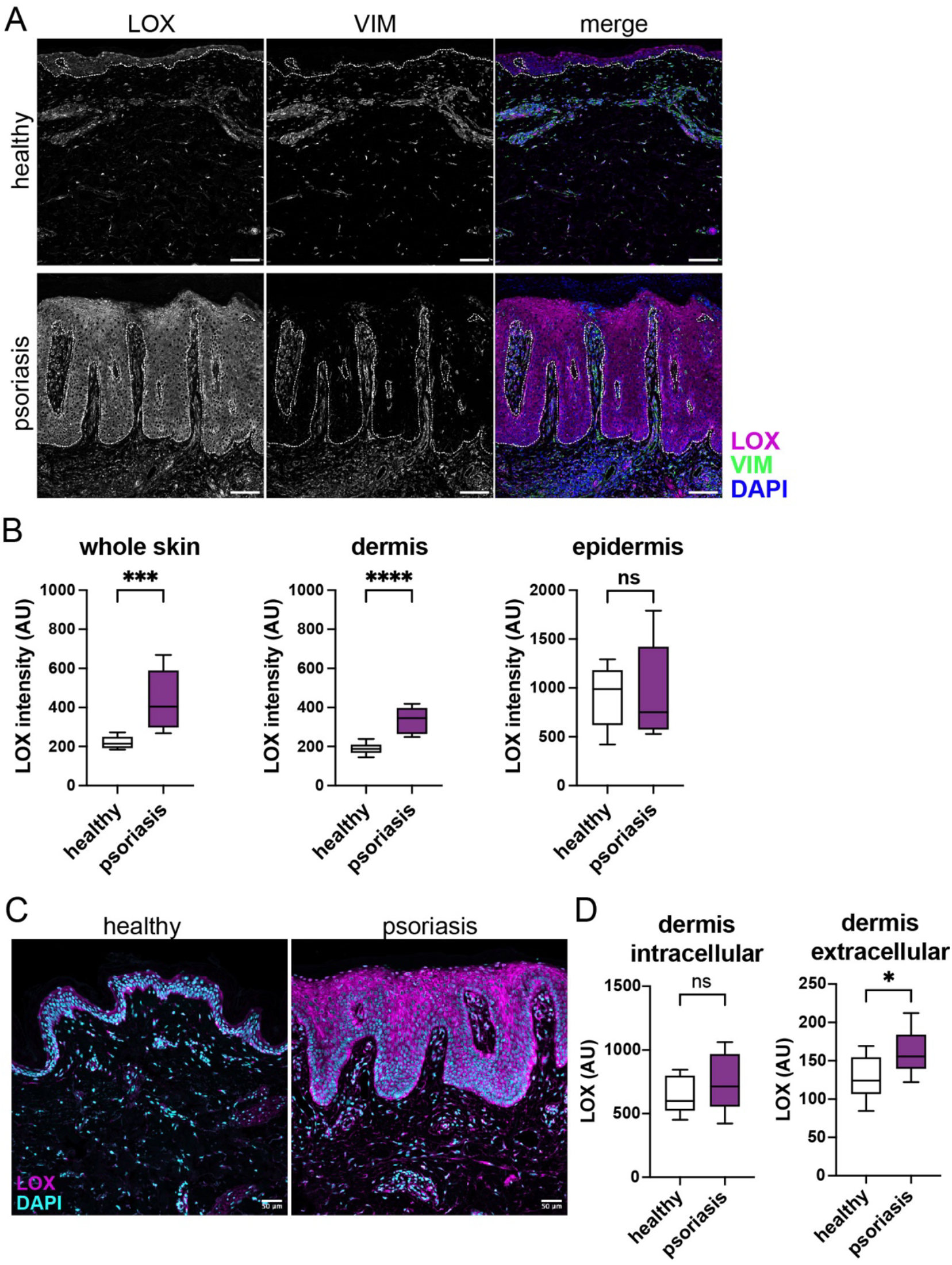
The ECM modifying enzyme LOX is highly expressed in psoriasis. **A)** Immunofluorescence staining of healthy and psoriatic skin sections is shown for LOX and VIM, and the merge with LOX in magenta, VIM in green and DAPI in blue. The fine dashed line indicates the epidermal-dermal border. Shown is one representative image for each. Scale bar = 100 μm. **B)** LOX intensities were measured in whole skin sections, the dermis and the epidermis of healthy (n = 11) and psoriatic skin (n = 8). **C)** Immunofluorescence staining of LOX in healthy and psoriasis skin. LOX is illustrated in magenta and nuclei in cyan (DAPI). Scale bar = 50 μm. **D)** Intracellular and extracellular LOX values were determined within the dermis of healthy (n=11) and psoriasis skin samples (n=8) using immunofluorescence staining and our ImageJ analysis pipeline.

**Figure 4.**
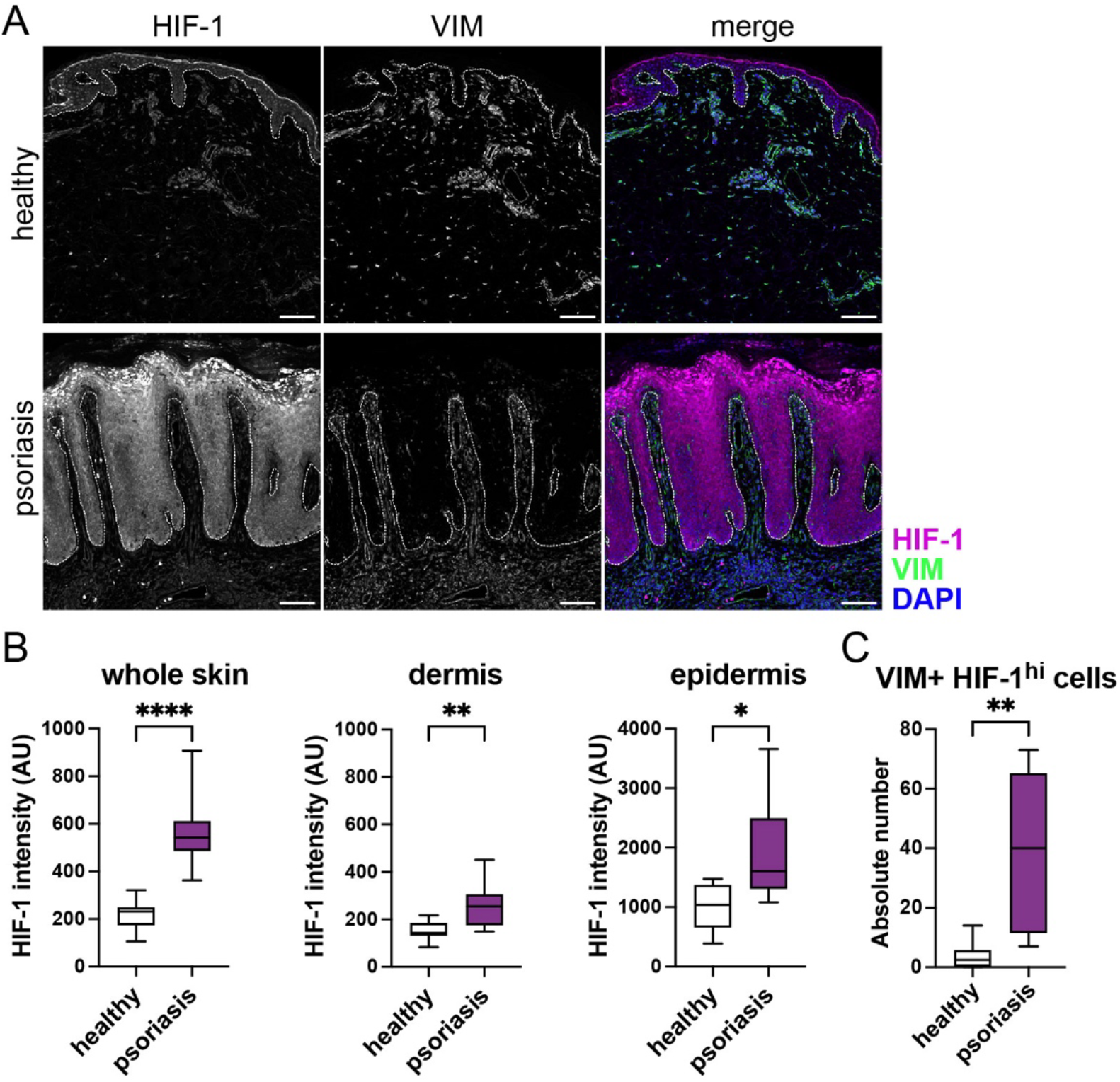
HIF-1 is upregulated in psoriasis. **A)** Immunofluorescence staining of healthy and psoriatic skin sections is shown for HIF-1 and VIM, and the merge with HIF-1 in magenta, VIM in green and DAPI in blue. The fine dashed line indicates the epidermal-dermal border. Shown is one representative image for each. Scale bar = 100 μm. **B)** HIF-1 intensities were measured in whole skin sections, the dermis and the epidermis of healthy (n = 9) and psoriatic skin (n = 9). **C)** VIM expressing cells were analyzed for HIF-1 expression. Shown is the absolute number of VIM+HIF-1^hi^ expressing cells in the dermis for psoriasis (n=8) and healthy skin (n=8).

Fibroblasts are the main producer of LOX within the dermis. We therefore co-stained skin sections for the mesenchymal marker vimentin (VIM) expressed by fibroblasts (Fig. 3A, middle panel) [22]. Single-cell data were extracted from images using our skin segmentation analysis tool, and LOX+ and VIM+ cells were plotted on scatter plots (Suppl. Fig. 3A). The number of VIM+ cells was significantly increased in the dermis of psoriasis patients compared to healthy skin (Suppl. Fig 3A and Suppl. Fig. 3B, left plot), which is in accordance with previous reports [23]. Further, LOX+VIM+ cells were elevated in psoriasis, although not significantly (Suppl. Fig. 3B, right plot).

To further investigate LOX expression on a single cell level, we revisited gene expression data from psoriasis and healthy skin and analyzed scRNA-seq data with a focus on fibroblasts (Suppl. Fig. 3C). We identified four different fibroblast clusters, which were extracted for detailed phenotypical analysis. Gene expression analysis for several ECM-relevant genes revealed that one particular fibroblast cluster (Cluster 13) expressed high amounts of LOX (Suppl. Fig. 3D). These fibroblasts were also expressing increased levels of the Microfibril Associated Protein 5 (MFAP5) and the Matrilin 4 (MATN4) gene, which both are known to be involved in ECM organization. Further, LOX-expressing fibroblasts also co-expressed pro-fibrotic markers, such as the Dipeptidyl Peptidase 4 (DPP4) and genes involved in ECM modifications, such as Procollagen C-Endopeptidase Enhancer 2 (PCOLCE) and the Bone Morphogenetic Protein (BMP1), which cleaves the LOX precursor to generate active LOX.

LOX needs to be secreted to perform its action. Previous studies reported that extracellular LOX attaches to ECM fibers within skin [13]. Thus, we separated epidermis and dermis *in silico*, masked all nucleated cells and quantified intra- and extracellular LOX intensities specifically in the dermis of healthy and psoriasis skin samples (Fig. 3C and 3D). Extracellular LOX was significantly increased in psoriasis compared to healthy skin, while intracellular LOX was marginally, but not significantly elevated (Fig. 3D). Our findings suggest the presence of a specific LOX-expressing fibroblast cluster which may play an important role in reorganizing the ECM in psoriasis.

### HIF-1 controls LOX secretion

When analyzing differential gene expression in cells belonging to Cluster 13, we found that more than 60% of fibroblasts from this cluster expressed the Hypoxia Inducible Factor 1 alpha (HIF-1), whereas only 40% of fibroblasts from healthy skin expressed this transcription factor (Suppl. Fig. 3E). As LOX expression is under the control of HIF-1, we investigated HIF-1 expression in skin sections using immunofluorescence microscopy (Fig. 4A). Indeed, we found HIF-1 to be highly expressed in psoriatic skin. The detailed quantification of HIF-1 intensity in whole skin, the dermis and the epidermis revealed a significant increase of HIF-1 expression in both skin compartments (Fig. 4B). Further, we found that HIF-1 was particularly upregulated in VIM expressing cells within the dermis (Fig. 4C, Suppl. Fig. 4).

To test a potential coordinated regulation of HIF-1 and LOX in fibroblasts, we performed *in vitro* experiments using Cobalt(II)-chloride (CoCl2), which stabilizes HIF-1 alpha protein and thereby induces its translocation from the cytoplasm into the nucleus in an oxygen-independent manner (Fig. 5). Skin fibroblasts were isolated from healthy human dermis and cultured with varying concentrations of CoCl2. HIF-1 protein was visualized using immunofluorescent staining and confocal microscopy (Fig. 5A). Quantification of nuclear HIF-1 confirmed that CoCl2 could induce the dose-dependent translocation of HIF-1 into the nucleus in primary fibroblasts, whereas the negative control (solvent) had no effect (Fig. 5B, Suppl. Fig. 5A-D). When we analyzed LOX protein in cell lysates and culture supernatants using Western blot, we found that the amount of secreted LOX was increased upon CoCl2 treatment, but cellular LOX seemed mostly unaffected (Fig. 5C). Quantification confirmed that 200 μM CoCl2 significantly induced LOX secretion in skin fibroblasts, whereas the effect on cellular LOX was minor (Fig. 5D). This suggests that in fibroblasts, elevated amounts of LOX are not accumulated within cells, but rapidly secreted to keep cellular LOX levels mostly stable. To validate a direct effect of HIF-1 on LOX secretion, we silenced HIF-1 in skin fibroblasts using small interfering RNA (siRNA). Both control cells (siCTR) and HIF-1 silenced fibroblasts (siHIF-1) were cultured with or without CoCl2, and nuclear HIF-1 intensities were measured to confirm the knock-down (Fig. 5E). In the absence of HIF-1, LOX secretion was not upregulated by CoCl2 (Fig. 5F and 5G). Our data suggest that HIF-1 regulates LOX secretion in human skin fibroblasts.

**Figure 5.**
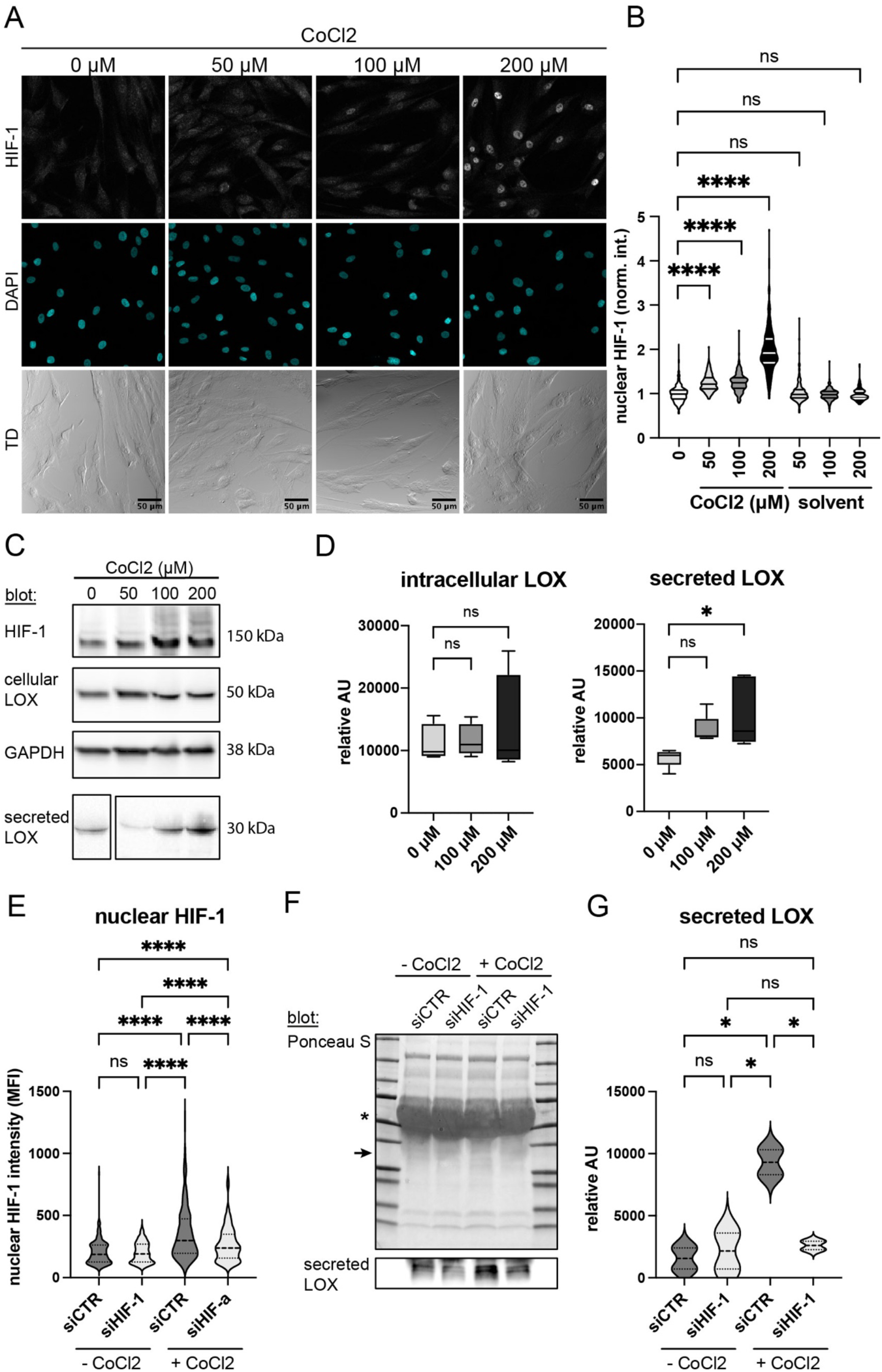
Chemical stabilization of HIF-1 induces LOX secretion. **A)** Skin fibroblasts were exposed to increasing concentrations of CoCl2 (50, 100, 200 μM), which prevents HIF-1 degradation, or the solvent alone (0 μM). Cells were fixed and stained for HIF-1 (upper panel) and DAPI (middle panel) to quantify nuclear HIF-1. The lower panel shows TD images of a representative experiment, scale bar = 50 μm. Statistical differences between groups were analyzed using the Fisher’s LSD test. **B)** Nuclear HIF-1 intensities were quantified from images of skin fibroblasts isolated from healthy skin upon culture with CoCl2 or the solvent. Shown is the normalized intensity of seven skin donors. **C)** Cell lysates of fibroblasts cultured with the indicated concentration of CoCl2 were analyzed for HIF-1 and cellular LOX using Western blot. GAPDH served as loading control. In addition, cell culture supernatants were analyzed for the presence of the secreted truncated form of LOX (30 kDa). **D)** Western blots from CoCl2 experiments (as described in C) were quantified for intracellular and secreted LOX (shown are data from fibroblasts derived from three skin donors and three independent *in vitro* experiments). **E)** HIF-1 was silenced in primary fibroblasts using siRNA (siHIF-1) and cells were exposed to 200 μM CoCl2 (+CoCl2) or the solvent (-CoCl2). Scrambled RNA sequences served as control for the RNA interference experiment (siCTR). Nuclear HIF-1 intensity was measured from immunofluorescence images as shown for A and B (shown are data from two independent experiments). **F)** Secreted LOX was analyzed in siHIF-1 and siCTR fibroblasts cultured with and without CoCl2. Ponceau S visualizes all proteins in cell culture supernatants illustrating equal loading. The big bands around 50-60 kDa represent serum from medium (marked with an asterisks). Secreted LOX was detected via Western blotting at 30 kDa (position indicated with an arrow). **G)** Secreted LOX in siCTR and siHIF-1 fibroblasts was quantified in supernatants of cells cultured without or with CoCl2 (200 μM) using Western blot (shown are data from two independent experiments).

### Substrate stiffness regulates LOX and HIF-1 in mechanosensitive skin fibroblasts

It is well established that LOX expression and activity is upregulated in fibrotic diseases resulting in changes in physical parameters of the ECM, such as altered stiffness, or rigidity ultimately affecting force propagation from the ECM to tissue-resident cells. Within skin, ECM stiffness varies depending on the site and age. ECM stiffness ranges from 5 kDa to 9 kDa and increases up to 30 kDa in fibrotic skin (Fig. 6A). To investigate whether fibroblasts are able to sense these physical changes, we cultured them on substrates with different stiffness ranging from 1.5 kDa to 28 kDa and glass, which has a stiffness of >1 GPa. Cells were fixed and stained for F-actin using the dye Phalloidin to explore force propagation from the substrate to fibroblasts (Fig. 6B). Stiffer substrates should thus induce stiffer cells, which would result in increased F-actin staining. Indeed, we found fibroblasts to be mechanosensitive, as F-actin assembly was upregulated in cells on stiffer substrate. In addition, nuclei changed in size and became bigger on stiffer substrate, further indicating mechanosensitivity of fibroblasts and F-actin regulation (Fig. 6B). Quantification of Phalloidin staining revealed a stiffness-dependent response, with a significant induction of F-actin from 1.5 kDa to 28 kDa and from 1.5 kDa to glass (Fig. 6C). When we investigated whether stiffer substrates may also affect LOX expression and/or secretion, we found that cellular and secreted LOX was increased in fibroblasts growing in a fibrotic environment (32 kDa) compared to fibroblasts on substrate with stiffness that resembled a “healthy” state (Fig. 6D). Plating fibroblasts on substrates with diverse stiffness raging from 2 kPa to 32 kPa revealed a significant stiffness-dependent increase of cellular and secreted LOX in fibroblasts (Fig. 6E). In a psoriasis mouse model it was reported that elevated IL-17 levels promote collagen deposition and induce collagen modifications via LOX [24]. We therefore wanted to investigate whether IL-17 has also an impact on LOX expression and/or secretion in human skin. However, incubation of fibroblasts with IL-17 did neither increase the amount of cellular or secreted LOX nor HIF-1 protein in fibroblasts (Suppl. Fig. 5F), suggesting that mechanical regulation is the main mechanistic driver in our system.

**Figure 6.**
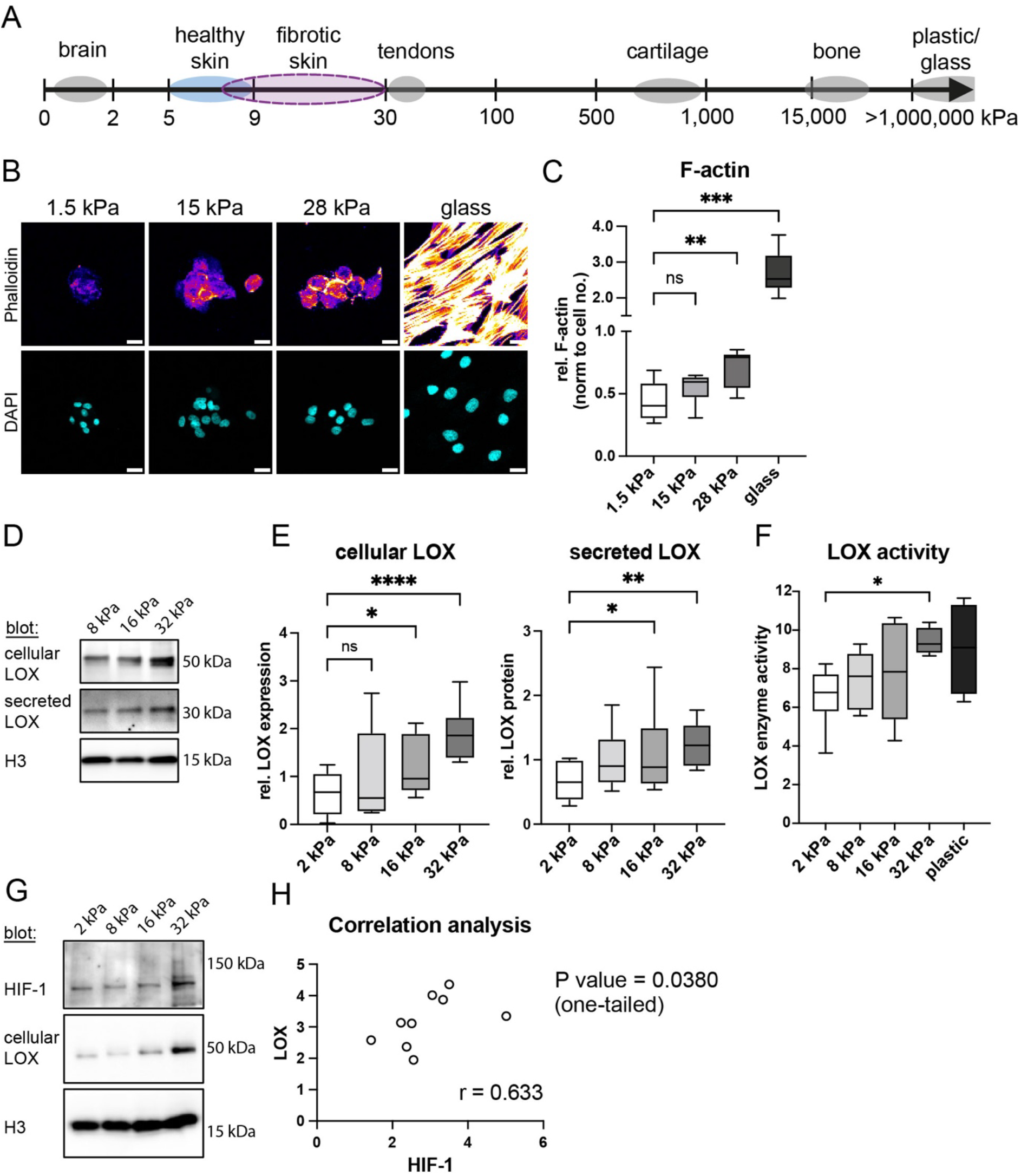
LOX and HIF-1 expression and LOX activity is increased in mechanosensitive skin fibroblasts in response to stiff substrate. **A)** Schematics of stiffness values in various tissues of the human body (in kPa). Common values for healthy (blue) and fibrotic skin (purple) are highlighted. **B)** F-actin cytoskeleton content was evaluated in fibroblasts that were cultured on substrates with various stiffness using Phalloidin. Nuclei were stained with DAPI and images were captured at a confocal microscope. Phalloidin is shown with the LUT Fire (high intensity in yellow/while and low intensity in blue/purple). **C)** F-actin was quantified in fibroblasts by measuring Phalloidin intensity and normalizing the intensity to the cell count (n=6). **D)** Cell lysates and cell culture supernatants of primary skin fibroblasts seeded on substrate with various stiffness ranging from healthy (8 kPa) to fibrotic skin (16 and 32 kPa) were analyzed by Western blot for cellular and secreted LOX. Histon H3 served as loading control for cell lysates. **E)** Skin fibroblasts were seeded on substrate of variable stiffness as shown in B and lysates and supernatants were analyzed for cellular and secreted LOX, respectively. Shown are extracted intensity values from Western blot analysis that were normalized to the mean of 8 kDa corresponding to the stiffness of healthy skin. Statistical differences were evaluated using matched 1-way ANOVA and the Fisher’s LSD test (n=6). **F)** LOX enzyme activity was measured in supernatants from skin fibroblasts seeded on substrate with various stiffness. Statistical differences were evaluated using matched 1-way ANOVA and the Fisher’s LSD test (n=6). **G)** HIF-1 protein expression was investigated in cell lysates of skin fibroblasts seeded on substrates with various stiffness by Western blot. Cellular LOX and H3 served as controls. **H)** LOX and HIF-1 mRNA was quantified using qPCR in skin fibroblasts isolated from different donors. Shown is the Spearman correlation analysis between HIF-1 and LOX (n=9).

As mentioned earlier, the 50 kDa LOX precursor is secreted into the extracellular space and cleaved into an active form with 30 kDa (Fig. 5C, Fig. 6D, Suppl. Fig. 5A). To verify that secreted LOX is functional in our *in vitro* system, we tested supernatants from fibroblasts grown on substrates with diverse stiffness for enzymatic activity and found a significant increase in LOX enzyme activity derived from fibroblasts grown on stiff substrate (Fig. 6F and Suppl. Fig. 6B). Our *in vitro* data firstly show that fibroblasts express both LOX and the respective proteinase, such as BMP1, responsible for cutting LOX into the catalytically active form, rendering them independent from other cells in this respect. Interestingly, also HIF-1 protein levels were increased in fibroblasts growing on stiff substrate compared to cells on soft substrate. We observed a significant correlation between HIF-1 and LOX expression (Fig. 6H). Our data uncover a yet unknown mechanism of mechanical induction of LOX secretion and activity in dermal fibroblasts under the control of HIF-1, which may fuel disease progression by changing the ECM architecture and tissue mechanics in psoriasis.

## Discussion

Our study reveals a potential new player contributing to psoriasis pathogenesis, the ECM modifying molecule LOX. We discovered substantial architectural changes to the ECM in the papillary dermis potentially causing aberrant skin tissue mechanics. Fibroblasts are the main producers and modulators of the ECM within the dermis, and it has been shown that papillary fibroblasts are crucial for ECM remodeling in wound healing [25, 26]. A recent study has proposed that a switch of fibroblasts towards a pro-inflammatory phenotype may amplify inflammation in psoriasis [23]. Extending the findings of this study, our data suggest that not only pro-inflammatory fibroblasts, but also ECM modulating subsets, such as LOX expressing fibroblasts reported here, play a role in disease progression. We observed a distinct reorganization of the ECM in the papillary dermis in psoriasis from a poorly oriented network-like mesh, which is typical for healthy skin [25], to highly aligned, thickened and elongated collagen fibers that were oriented perpendicular to the skin surface. The decrease in ECM fiber entropy and increase in fiber alignment and differential orientation may affect skin tissue mechanics potentially influencing psoriasis pathogenesis on multiple layers: 1.) by guiding immune cells towards the epidermis, 2.) controlling vasculature modifications, and 3.) altering the force load within skin tissue and its propagation to tissue resident cells.

LOX is one of the main ECM crosslinking enzymes. Only 20% of ECM modifications are performed by other isoforms, such as the LOX-like enzyme LOXL1, which is mainly involved in elastin crosslinking [27]. LOX knock-out mice are not viable and display shorter and reduced elastic fibers and loosely dispersed unbundled collagen within skin, highlighting the essential role of this enzyme in development and skin integrity [27-29]. Within the dermis VIM+ fibroblasts regulate tissue modifications during homeostatic regeneration and upon wounding [13, 22, 23, 30]. We found an increase of VIM+ fibroblasts in psoriatic skin and identified a distinct fibroblast subset that produces LOX. Further, we discovered that the endopeptidase BMP1, the main producer of active LOX [16], is upregulated in psoriasis. These data suggest that fibroblasts specialized in ECM modifications are highly abundant in active lesions of psoriasis patients potentially fuel mechanical disturbances in tissue homeostasis.

Hypoxia is induced within psoriatic lesions by increased metabolic activity, hyperproliferation of skin resident cells and infiltrating immune cells. We and others showed that HIF-1 expression is upregulated in the skin of psoriasis patients [6, 7, 31]. It has been reported that LOX expression is regulated via the transcription factor HIF-1 [15], but a linkage between HIF-1 stability and LOX secretion in fibroblasts has not been shown so far. Our data suggest that LOX secretion by fibroblasts is regulated in a HIF-1-dependent, hypoxia-independent manner through the sensing of and responding to physical parameters, such as ECM stiffness, which also directly affected the F-actin cytoskeleton, confirming mechanosensitivity. Further, we discovered that HIF-1 can be upregulated through mechanical forces exerted on fibroblasts, again independent of hypoxia. A recent study investigated mechanisms for injury-induced HIF-1 expression in epithelia using a mouse model and found the hypoxia-independent induction of HIF-1 through IL-17 [32]. This is in contrast to our study, reporting that IL-17 did not play a role, at least in cultured fibroblasts, suggesting that they may contribute to tissue structure maintenance and integrity in the absence of this cytokine. Additionally, the finding that LOX and HIF-1 expression is strongly induced in wound healing [16] supports the idea of a mechanical regulation of LOX secretion. Interestingly, elevated levels of circulating LOX have been correlated with compromised cardiac function due to heart tissue stiffening [29]. Increased LOX secretion and serum levels may hence adversely affect cardiovascular, rheumatological and respiratory comorbidities observed in psoriasis patients. Similar findings have been reported for the cartilage oligomeric matrix protein COMP, which is elevated in the serum of both psoriasis and psoriatic arthritis patients and may drive both diseases [33]. However, further research is needed to understand the role of soluble ECM molecules in psoriasis comorbidities.

The observed mechanical induction of HIF-1 stabilization and LOX secretion in fibroblasts may create a positive feedback loop in psoriasis that could adversely affect disease clearance. Furthermore, this may also initiate lesions after physical trauma or mechanical stress in non-lesional skin, a phenomenon known as Koebner effect [9]. We identified several ECM-relevant genes that were differentially regulated in non-lesional skin, such as various collagens. Single nucleotide polymorphisms in the genes COL6A5, which is involved in cell adhesion, and COL8A1, which plays a role in blood vessel formation, have been linked to a higher prevalence of psoriasis [34]. We found these genes to be differentially regulated in non-lesional psoriatic compared to healthy skin. Additionally, we found several genes that are important for stabilizing the epidermal-dermal junction (COL4A6, COL7A1), involved in ECM modifications (MMP7) or cell-matrix interactions (THBS1, ICAM1) to be differentially regulated in non-lesional versus healthy skin [10]. Hence, one may speculate that non-inflamed skin from psoriasis patients differs structurally and mechanically from healthy skin and thereby provides an ideal basis for lesion initiation. Indeed, Dobrev *et al*. found that the mechanical properties of NL skin differed significantly from those of normal healthy skin, showing less distensibility and higher viscoelasticity when pulled, which was independent of epidermal hyperplasia [5]. This suggests that the dermis, in particular the papillary dermis and the collagen structure within, may affect tissue mechanics even in non-inflamed skin [5]. Further, Buehler *et al*. showed that the length of collagen fibrils directly correlates with higher stability [12]. We found longer and thicker collagen fibers in the papillary dermis, which would explain a resistance to pulling forces and perpendicular orientation of fibers may further enhance this effect. However, more research is needed to generate a detailed understanding of the role of ECM modifications in disease initiation.

One of the hallmarks of psoriasis is the influx of inflammatory immune cell subsets into the skin, such as T cells, neutrophils, macrophages, and dendritic cells [3]. The dermis is the place where most of the immune cells reside and through which cells need to migrate to reach the epidermis. Hence mechanical perturbations of the interstitial matrix can be sensed by tissue resident cells and transformed into altered biochemical signals or have a tremendous impact on immune cells regarding cell adhesion, migration, proliferation and differentiation [8, 11, 35, 36]. For instance, the perpendicular orientation of collagen fibers in psoriasis may create migration tracks that guide immune cells directly to the epidermis. CD4+ helper T cells associate and move along or in between fibronectin or chemokine coated collagen fibers and bundles within the dermis [37]. Collagen crosslinking can also modulate the topology of collagen fibers creating rough surfaces, which can be used by migrating T cells to generate friction between them and the ECM for efficient 3D migration without depending on focal adhesions [10, 38, 39]. Substrate stiffness can further regulate the switch between amoeboid friction-based movement and mesenchymal adhesion-based traction force migration suggesting that the modified ECM structure in psoriatic skin may also affect the mode of migration of T cells and enhance their motility [40]. Highly aligned collagen fibers, as seen in psoriasis, can also affect the directionality and speed of T cells. This has been shown for CD8+ T cells, which move faster and more directed in aligned collagen compared to unstructured collagen while having a rounded morphology with low protrusions suggesting the adaptation of an amoeboid migration mode along collagen fibers [41]. Hence, restoring the papillary ECM to a healthy state, which is more parallel to the epidermis could block CD8+ T cell migration into the epidermis and thereby prevent disease development [3]. On the contrary, highly fibrotic dense ECM structures can also prevent T cell migration and limit cytokine diffusion [23, 42]. Tumors are often encapsulated by highly crosslinked ECM structures thereby preventing T cell access. Inhibition of LOX crosslinking has been shown to reduce the stiffness of tumor-surrounding matrix and increased T cell motility in a mouse tumor model [42]. We further found that CD45+ immune cells accumulated within dermal pockets and realigned collagen fibers built the structural scaffold. By this, disease driving T cells that produce IL-17 or that support papillary fibroblast survival may be kept in the skin [3, 43]. Ma *et al*. showed that proinflammatory cytokines are trapped in the tips of dermal papillae potentially directing cell migration towards them [23]. ECM modifications could play a determining role in this process as dense matrix can limit cytokine diffusion and create chemokine gradients by sequestering soluble molecules in addition to limiting tissue cell migration [10]. Hence, ECM modifications in psoriasis may on the one hand promote and direct immune cell migration and on the other hand prevent immune cell evasion out of the dermis, keeping up a proinflammatory milieu further fueling the chronicity of psoriasis.

In conclusion, LOX may represent a novel therapeutic target to specifically tackle ECM modifications in psoriasis. Integrating our insights into ECM modifications with established pathways involved in psoriasis pathogenesis could offer new adjunctive treatment avenues to enhance the long-term effectiveness of existing therapies. Indeed, glucocorticoids have been shown to soften the interstitial matrix, potentially providing additional “beneficial” side-effects although not targeted [44]. Currently, we do not yet fully understand the mechanisms behind stiffness-dependent HIF-1 stabilization and LOX secretion in skin fibroblasts. Future studies will be needed to shed light onto novel roles of HIF-1 regarding LOX regulation and will open promising new options to target LOX in psoriasis and beyond.

## Material & Methods

### Skin tissue and ethics declaration

Healthy human skin samples were sourced from excess skin, which was removed in the course of unrelated cosmetic surgery after written informed consent. Ethical approval for the study was obtained from the Ethics Committee of the Medical University of Vienna. Before processing, skin was washed briefly in 70% ethanol and PBS. Biopsies were then taken for dermal cell isolation and paraffin embedding. Formalin-fixed paraffin-embedded (FFPE) tissue blocks were prepared after skin fixation in 7.5% formalin for 24 hours and further processed using a Tissue-Tek VIP® 6 AI-Tissue processor. Psoriasis samples were sourced from FFPE blocks of skin biopsies originally taken for clinical routine diagnostics from patients with psoriasis at the Department of Dermatology, Vienna General Hospital. Diagnostic workup of this material has been finished, with FFPE blocks stored in the routine laboratory as retained archival material. Secondary use of the material was approved by the Ethics Committee of the Medical University of Vienna (#1173/2021), where laboratory staff ensured that after sampling sufficient tissue remained for potential future clinical use.

### Isolation of dermal cells, expansion and culture

For dermal cell isolation, human skin was cut into thin pieces and incubated overnight at 4°C with dispase (Roche) at 2.4 U/ml. Next day the epidermis was separated from the dermis with two forceps [45]. The dermis was further cut up and digested with liberase (Roche) at 37°C for 90 minutes. Cell culture media was added, the cell suspension filtered through a 100 μm and 70 μm filter, and centrifuged at 350 g for 7 minutes. The resulting pellet was resuspended and cultured in fibroblast medium (DMEM, 10% FCS, 1% Penicillin/Streptomycin, and 1% L-Glutamine) at 37°C and 5% CO2. Media changes were conducted every 3 days, and cells splitted at 80% confluency using 0.05% Trypsin-EDTA.

### CoCl2 assay

A total of 5,000 primary fibroblasts isolated from healthy donors were seeded per well into an 8-well slide (IBIDI). The medium in each well consisted of 250 μL of DMEM (Gibco) supplemented with 10% FCS, 1% Pen/Strep, and 1% L-Glutamine. The cells were allowed to adhere overnight at 37°C. Cobalt (II) chloride hexahydrate (CoCl2) (Sigma-Aldrich) was dissolved in water to achieve a 25 mM solution, which was purified by filtration through a 0.22 μm filter to ensure sterility before use. The following day after seeding the fibroblasts, CoCl2 was added to the cells at concentrations of 50 μM, 100 μM, and 200 μM. Sterile water was used as the vehicle control for the experiment. Additionally, a control consisting of simple DMEM without any CoCl2 or water was included. After 24 hours, the CoCl2 treated cells were used for further experiments.

### siRNA knock-down

Lyophilized ON-TARGET siRNA directed against human HIF-1A mRNA (J-004018-07-0002) or a non-targeting control (non-targeting siRNA #1) were purchased from Dharmacon. siRNA tubes were briefly spun down, siRNA resuspended in PCR-grade water according to the manufacturer’s instructions, aliquoted and stored at -80°C. For siRNA knock-down experiments, skin fibroblasts were seeded at a density of 1.5 x10^4^ cells/cm^2^, medium was changed the next day to Opti-MEM supplemented with 1% Pen/Strep, and 1% L-Glutamine (Gibco) and cells transduced using 50 nM siRNA and DharmaFECT 1 (Dharmacon) according to the manufacturer’s protocol. After a 5 hour incubation at 37°C, medium was exchanged to fresh Opti-MEM. The following day, cell lysates were harvested or cells were further treated with 200 μM CoCl2 or the solvent. In this case cell lysates and supernatants were harvested for Western blot analysis after culturing the cells for another 24 hours.

### Preparation of isolated skin cells for imaging

For fluorescent imaging of primary fibroblasts, cells were cultured in imaging dishes and washed with 1X PBS before fixation using 4% paraformaldehyde (PFA) for 15 min at RT. Cells were washed once with 1X PBS and permeabilized using 0.1% Triton X-100 for 15 min at RT. For HIF-1 staining, cells were incubated with unconjugated rabbit anti-human HIF-1 monoclonal antibody (mAb, Abcam, ab51608) or the corresponding rabbit isotype control Ab (Abcam, ab172730) using a concentration of 10 μg/mL for 1 hour at room temperature. Cells were washed three times with 1X PBS before incubation with secondary goat-anti-rabbit Alexa Fluor 568 Ab (Invitrogen) for 30 min at RT. For staining of F-actin, cells were incubated with AF647-labelled Phalloidin (Invitrogen). Cells were washed three times with 1X PBS before nuclei were stained with 4′,6-diamidino-2-phenylindole (DAPI, Roche) for 10 min at RT. Before imaging, cells were washed once more and overlayed with 1X PBS.

### Immunofluorescence staining of skin tissue sections

For immunofluorescence staining of FFPE samples, skin tissue was sectioned with 4 micron thickness, and deparaffinization was conducted by melting the paraffin followed by rehydration through multiple immersions in xylene and descending concentrations of ethanol. Antigen retrieval was conducted using citrate buffer (pH 6) in a pressure cooker (2100 Retriever). After cooling to room temperature, slides were washed thrice with 0.3% Triton X-100 in PBS (Sigma) and once with PBS before staining. The following primary (anti-human) and secondary antibodies were used: anti-LOX (Abcam, ab174316), anti-vimentin (Dako, M7020), anti-E-cadherin (Abcam, ab1416), anti-HIF-1 (Abcam, ab51608), anti-CD45 (Atlas Antibodies, AMAb90518), and goat anti-mouse IgG2a cross-adsorbed secondary Ab, Alexa Fluor™ 647 (Thermo Fisher Scientific, A-21241), goat anti-mouse IgG1 cross-adsorbed secondary Ab, Alexa Fluor™ 488 (Thermo Fisher Scientific, A-21121), goat anti-rabbit IgG (H+L) cross-adsorbed secondary Ab, Alexa Fluor™ 568 (Thermo Fisher Scientific, A-11011), goat anti-mouse IgG1 cross-adsorbed secondary Ab, Alexa Fluor™ 647 (Thermo Fisher Scientific, A-21240). Briefly, primary antibodies were diluted in 2 % BSA-PBS, applied to tissue sections and incubated over night at 4°C. After washing, secondary antibodies plus 10% goat serum were applied and incubated for one hour at room temperature, before washing and nuclei staining with DAPI. DAPI was not used for CD45-stained slides due to interference with SHG imaging. Slides were mounted with fluorescence mounting medium (Dako).

### Masson-Trichome staining

Masson-Trichome staining (Roth) was performed according to the manufacturers protocol with some modifications. Briefly, paraffin-embedded tissue slices were deparaffinized, rehydrated and incubated in preheated Bouin’s solution at 56°C for 15 minutes, followed by a brief rinse with water for 1-2 minutes. Then, nuclei were stained in Weigert’s Iron Hematoxylin for 10 minutes, followed by thorough rinsing with distilled water for 10 minutes. Slides were then immersed in Goldner’s stain 1 for 10 minutes, rinsed with 1% acetic acid for 30 seconds and placed in Goldner’s stain 2 for 5 minutes followed by a 30 second rinse 1% acetic acid. Finally, slides were placed in Anillin Blue staining solution for 5 minutes, and then incubated in 1% acetic acid for 2 minutes. To complete the staining, dehydration steps were performed by placing the slides in 75%, 85%, and 100% ethanol for one minute each, followed by air-drying. Stained slides were mounted using Eukitt mounting media and visualized using a brightfield/epifluorescence microscope (Leica DM6000B) and a 10x objective.

### Confocal and Multiphoton microscopy

For imaging of skin sections we used a confocal laser scanning microscope (FV3000 on inverted IX83-SSU, Olympus). The microscope was equipped with a 10x objective (NA 0.4), four laser lines (405 nm, 488 nm, 561 nm, 640 nm; Coherent), and a spectral detector. 1272 x 1272 micron images at a resolution of 0.3 μm/px were taken to capture both the epidermis and dermis of sectioned skin biopsies.

An upright multiphoton microscope (FV31S-SW, Olympus) was used for the label-free visualization of ECM fibers using second harmonic generation (SHG) microscopy. The microscope was equipped with a water-dipping 25x IR objective (NA 1.05), two tunable infrared laser lines (INSIGHT X3 and MaiTai, both Spectra-Physics), two PMT detectors and two GaAsP detectors, and 509 x 509 micron images were captured with a resolution of 0.5 μm/px. For some experiments CD45 staining was used to simultaneously visualize immune cells and the ECM via SHG. For this a dichroic mirror (SDM570) was used to detect CD45 labeled with AF647 between 660 and 750 nm and SHG between 426-477 nm using excitation pulsed laser wavelengths at 1200 nm and 880 nm respectively.

### Analysis of tissue sections

The segmentation macro SCAnED was used for the analysis of cell numbers and average protein expression levels across diverse skin cell populations. The macro was designed in our lab and runs on the free image analysis software Fiji [46]. Briefly, this macro aids in segmenting epidermal and dermal compartments in skin tissue and in identifying and quantifying cells and nuclei within these compartments. Particularly adept at handling multi-channel immunofluorescence images, it enables robust analysis of up to five distinct marker of interest channels. The expression of intracellular and extracellular/secreted LOX was measured within the dermis. For this the keratinocytes were stained with E-cadherin and used to mask and exclude the epidermis for analysis. Thereafter, the total mean intensity of intracellular LOX was determined in Fiji by creating a mask that covers all dermal cells. For extracellular/secreted LOX, the mask was inverted to measure LOX intensities within the dermis excluding cellular LOX.

### Analysis of ECM fibers

The MATLAB tool CurveAlign was used to quantify collagen fiber angles relative to the skin surface and CT-FIRE was used to segment and extract data from individual fibers from SHG images [20]. Overall fiber length and width was estimated in whole skin and dermal papillae using SHG images of skin sections and the ImageJ analysis tool Ridge Detection [47]. The following settings were used: Line width (4), high contrast (100), low contrast (40), sigma (1.65), lower threshold (1), upper threshold (2.72), minimum length (8).

### Analysis of nuclear translocation of HIF-1

Immunofluorescence staining of nuclear HIF-1 was quantified using FIJI. Images of DAPI-stained nuclei were appropriately thresholded to generate a mask. Subsequently, the image was converted to a binary format, and the watershed function was applied to enhance the accurate selection of nuclei. Particle analysis was conducted using a range of 50 to infinity pixels and circularity values between 0.3 and 1. The intensity of HIF-1 staining was then measured.

### Western blotting

Cell culture supernatants were collected, cOmplete protease inhibitor (Roche) was added and samples stored at -20°C until further analysis. Cells were washed with pre-cooled 1X PBS and lysed for 20 minutes on ice using RIPA buffer (Thermo Fisher scientific) containing protease inhibitor. Cell lysates were harvested and stored at -20°C. For Western blot analysis, supernatants and lysates were thawed on ice, centrifuged at 10 000 g for 5 min and mixed with 4x Laemmli buffer (Bio-Rad) and dithiothreitol (DTT), before heating to 95°C for 5 min. Proteins were separated via SDS-PAGE using precast 4-20% gradient gels (Bio-Rad, Criterion TGX) and blotted onto nitrocellulose membranes (supernatants) or PVDF membranes (cell lysates). After transfer, membranes were blocked with 5% bovine serum albumin (BSA) for 1 hour at room temperature. Proteins were detected using rabbit anti-HIF-1 (Novus, NB100-449), rabbit anti-LOX (Abcam, ab174316) and rabbit-anti histone H3 (Cell Signaling, #4499) mAbs in 5% BSA shaking overnight at 4°C, followed by washing and incubation with HRP-conjugated goat anti-rabbit IgG Ab (Bio-Rad) for 1 hour at room temperature. LOX and histone H3 were developed using the Clarity Western ECL substrate solutions (Bio-Rad), while HIF-1 was detected using high sensitivity Clarity Max Western ECL (Bio-Rad). For visualization of total protein content in supernatants, Ponceau S was used. Chemiluminescence and colorimetric images were captured using a ChemiDoc machine (Biorad). For re-probing of membranes, antibodies were removed using a mild stripping buffer (1.5% (w/v) glycine, 1% (w/v) SDS and 1% (v/v) Tween 20; pH adjusted to 2.2 using HCl). Membranes were thoroughly washed and blocked again with 5% BSA for 1 hour before re-probing.

### Fibroblast Stiffness Assay

To test whether fibroblasts are mechanoresponsive, 1.5 x 10^5^ cells were cultured on 35 mm dishes with substrates of varying stiffness. Briefly, CytoSoft 6-well plates, with stiffness ranging from 2, 8, 16 and 32 kPa (Advanced BioMatrix) or ESS μ-dishes (1.5, 15, 28 kPa Ibidi) were coated with rat-tail collagen I (Ibidi) at a final concentration of 100 μg/ml in PBS for one hour at room temperature. After washing with PBS, primary fibroblasts from healthy skin were seeded onto the substrates and left to adhere in DMEM+10%FCS. The next day, medium was changed to serum-free phenol-free OptiMEM and cells were cultured for another 24 hours before harvesting cell culture supernatants and cell lysates for LOX activity measurements, Western blot and qPCR.

### *In vitro* analysis of HIF-1 protein, LOX protein and activity

Enzymatic activity of secreted LOX was measured in 10 μl cell culture supernatants using the fluorometric Lysyl Oxidase Activity Assay Kit (Abcam). LOX activity kinetics was evaluated by measuring fluorescence (Ex/Em = 535/587 nm) at 37°C every 30 seconds for 90 minutes in non-binding, black 96-well plates (Corning) using a spectrophotometer.

### mRNA isolation and qPCR

Cellular RNA was isolated using TRIzol reagent (Invitrogen) according to the manufacturer-recommended protocol dissolved in RNase-free water before storage at -80°C. cDNA was prepared using the RevertAid First Strand cDNA Synthesis Kit (Thermos Fisher). For quantitative real-time PCR (qPCR), the Taq™ Universal SYBR® Green Supermix (BIO RAD, 1725121) was used. qPCR was performed on a Step One Plus cycler (Applied Biosystems) using the following primer (5’-3’): LOX_iso1_fwd: CACTCCGATCCTGCTGATCC, LOX_iso1_rev: TAGCCAGCTTGGAACCAGTG, HIF-1a_fwd: GCTTTAACTTTGCTGGCCCC, HIF-1a_rev: TTTTCGTTGGGTGAGGGGAG, GAPDH_fwd: AAGGTGAAGGTCGGAGTCAA, GAPDH_rev: AATGAAGGGGTCATTGATGG, ACTB_fwd: CGCCGCCAGCTCACC, ACTB_rev: CACGATGGAGGGGAAGACG. Data were analyzed using the ΔΔCt method.

### Single cell RNA sequencing (scRNA-seq) and Affymetrix gene expression analysis

scRNA-seq analysis was performed using published data from Gao *et al*. consisting of 3 healthy and 3 psoriasis skin samples [18]. The raw gene counts matrices were downloaded from GEO (accession no. GSE162183). Each sample underwent a quality control pipeline individually, before merging the datasets and analyzing them with Seurat [48] as described previously [49]. Microarray gene expression data from Yao *et al*. comprising 80 healthy (H), lesional (L) and non-lesional (NL) psoriasis skin samples (Affymetrix Human Genome U133 Plus 2.0 Array, GSE14905 [19]) were analyzed using Genevestigator (Immunai/NEBION). Over 700 differentially regulated genes were identified using the linear model for micro-array data (limma) framework comparing NL and H skin. Gene lists were further analyzed using the Reactome gene set enrichment online tool, which identified ECM organizing genes.

### Statistical analysis

Statistical tests were performed using unpaired parametric two-tailed Student’s t tests to compare the mean of two groups. Unless indicated otherwise, we used one-way ANOVA to compare the mean of multiple groups with each other, followed by Tukey or Bonferroni post-hoc tests using GraphPad Prism 10 (version 10.2.0). Statistical tests for gene expression analysis from single cell data were performed using the Wilcoxon rank sum test. Correlation analysis between HIF-1 and LOX expression was performed using the nonparametric Spearman correlation test with a 95% confidence interval (one-tailed). For statistical evaluation of stiffness assays analyzed with Western blot, matched samples were analyzed using one-way ANOVA or the Mixed-effects model with a Fisher’s LSD test. Overall P < 0.05 was considered statistically significant. Potential outliers were identified using the ROUT method.

## Acknowledgements

We would like to thank all patients for agreeing to provide material for research. Further, we would like to thank the Land Niederösterreich (Danube ARC) and the Fellinger Krebsforschung for funding our study, and the Vienna Scientific Cluster of the Technical University of Vienna for providing computing space at their HPC.

## Author contribution

PB, PW, DS performed experiments and analyzed data, NZ prepared skin sections, MB established staining protocols, WW provided critical input, MW supervised the statistical analysis and analysis of scRNA-seq data, KP conceived, planned and supervised the study, planned, performed and analyzed experiments and wrote the manuscript. All authors proof-read the manuscript.

## Supplementary Material

Supplementary Figures 1-6

