## Supplementary material for "Stiffness-dependent LOX regulation via HIF-1 drives extracellular matrix modifications in psoriasis": Supplementary Material LOX in psoriasis.pdf

### Supplementary Material – Balsini et al.

#### Supplementary Figure 1 - Balsini et al.

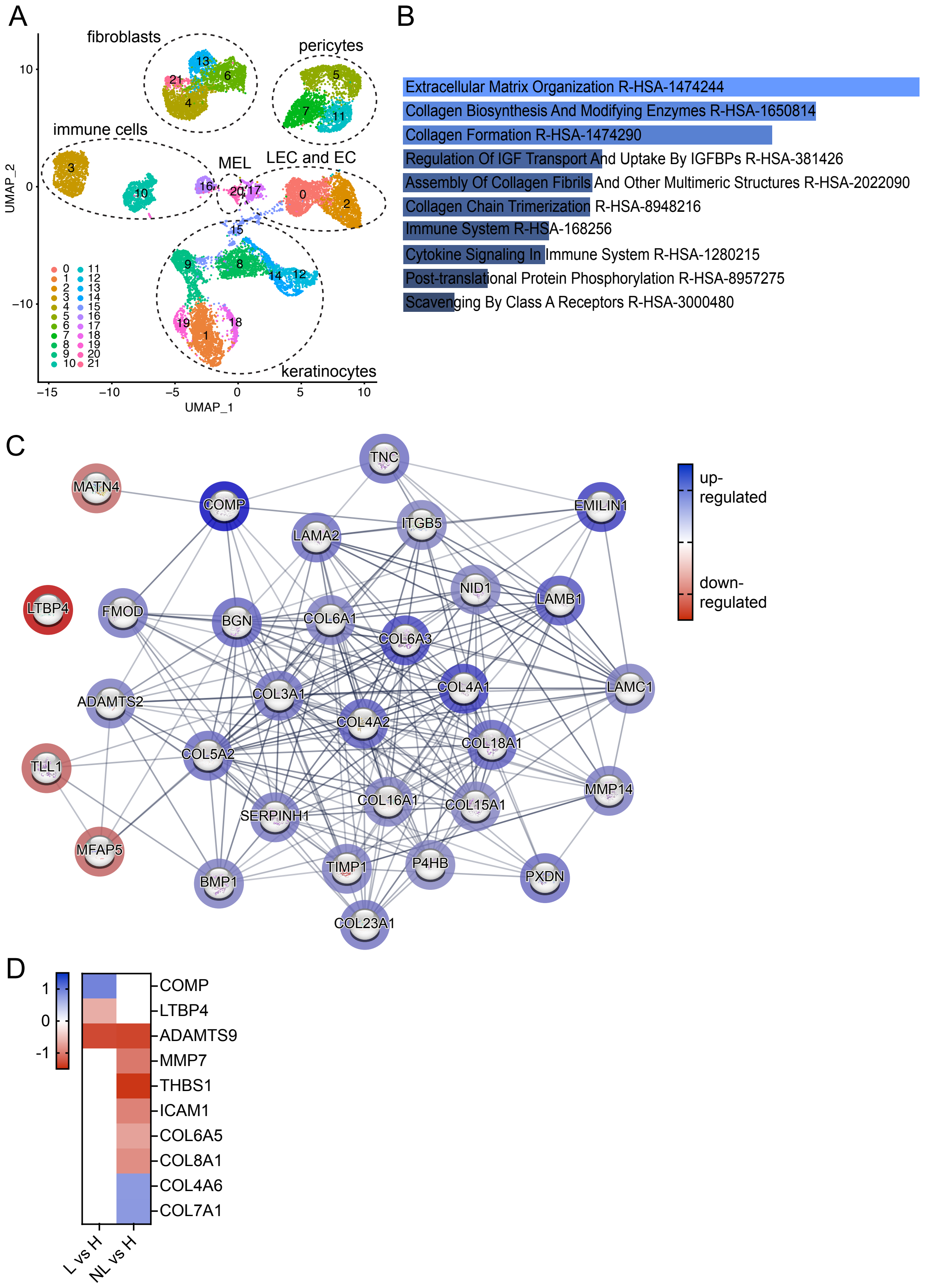

**Supplementary Figure 1. Extracellular matrix organization is differentially regulated in psoriasis compared to healthy skin. A)** Single cell RNA sequencing analysis of 3 healthy and 3 psoriasis skin biopsies. After quality control datasets were merged and 16,751 cells were analyzed using Seurat. Shown is a UMAP dimensionality reduction revealing 22 clusters corresponding to all known skin cell types. Fibroblasts, pericytes, immune cells, melanocytes (MEL), lymphatic endothelial (LEC) and endothelial cells (EC) and keratinocytes were highlighted. **B)** Differentially regulated genes from the comparison of psoriasis and healthy skin were analyzed with EnrichR. Shown are the top 10 enriched GO Biological Process 2023 pathways. **C)** Differentially regulated genes were further analyzed with Reactome and STRING. Illustrated are up- (blue) and down-regulated genes (red) from the identified Reactome pathway "Extracellular matrix organization" that were further analyzed with STRING to visualize putative interactions. **D)** Bulk gene expression data from lesional (L), non-lesional (NL) psoriatic and healthy (H) skin (80 samples, Affymetrix Human Genome U133 Plus 2.0 Array, GSE14905) were analyzed for ECM organization-relevant genes. Shown are fold change values for the comparisons L vs H and NL vs H with upregulated genes in blue and downregulated genes in red.

Supplementary Figure 2 - Balsini et al.

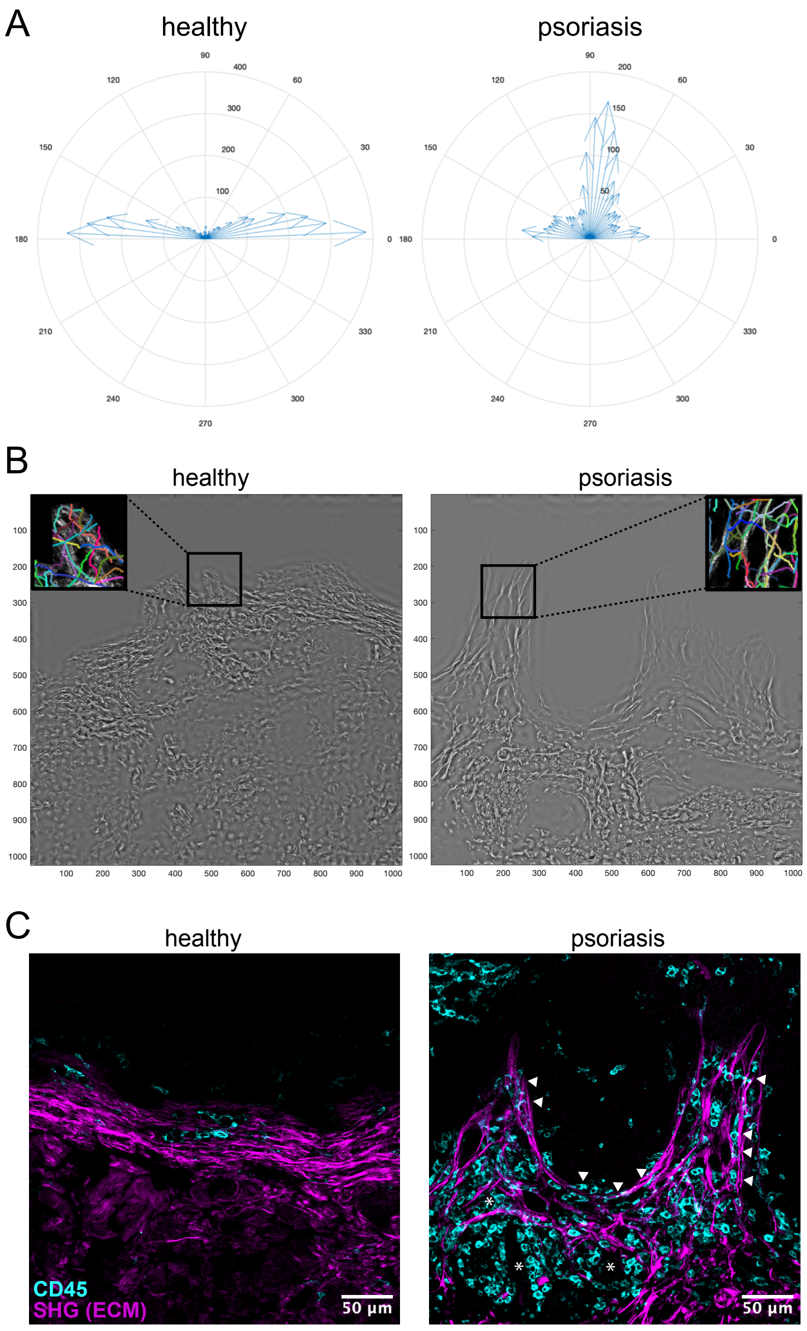


**Supplementary Figure 2. Quantification of ECM modifications in the dermis. A)** Compass plot of the angle of fibers analyzed in bulk using CurveAlign. **B)** Individual fibers were detected using CT-FIRE allowing subsequent determination of single fiber metrics. Shown are representative images for of reconstructed ECM in healthy and psoriasis skin (grey image). Insets depict identified fibers within dermal papillae colored individually. **C)** Heathy and psoriasis skin samples were stained with anti-CD45 Abs to detect immune cells (cyan) and the ECM (magenta) simultaneously using a multiphoton microscope. Immune cells align with collagen fibers in the upper dermis and dermal papillae. Representative images, scale bar = 50 μm.

Supplementary Figure 3 - Balsini et al.

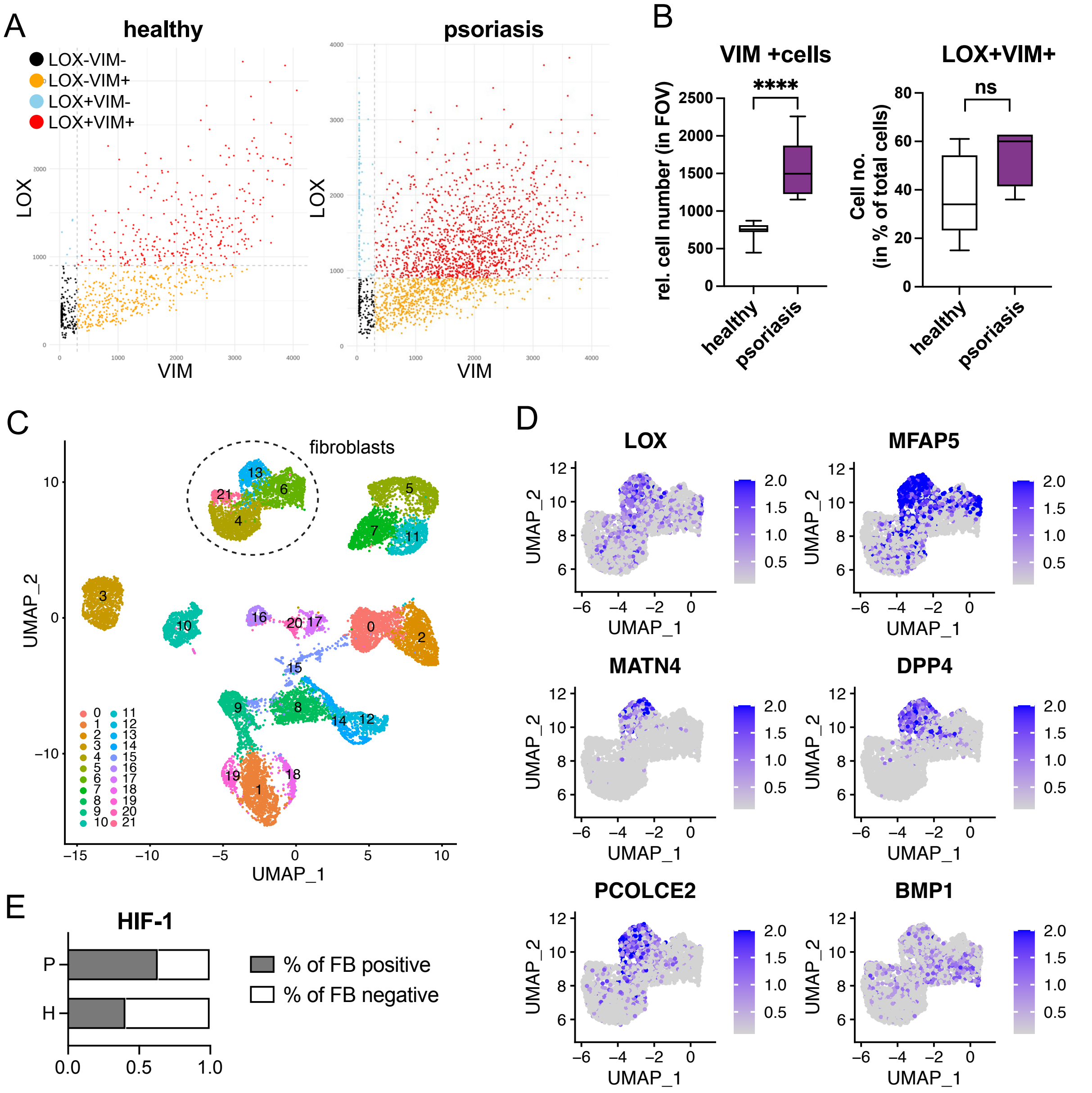


**Supplementary Figure 3. LOX is expressed by a distinct skin fibroblast subset. (A)** Expression analysis of dermal cells extracted from immunofluorescence images of healthy and psoriatic skin. Shown are representative dot plots illustrating LOX and VIM expression of extracted cells (each dot represents a cell). A gate was set to separate negative cells (black) from single positive (LOX+, blue; VIM+ orange) and double positive cells expressing both marker (LOX+VIM+, red). **B)** Data for VIM+ and LOX+VIM+ cells were extracted from immunofluorescence images of healthy and psoriasis skin using our image segmentation tool. Left plot: quantification of VIM+ cells in dermis. Shown is the relative cell number per field of view (n=9 for each). Right plot: the number of LOX+VIM+ cells in dermis is shown in percent relative to total cells for healthy (n=9) and psoriatic skin (n=4). **C)** UMAP plot with highlighted fibroblast subsets that were used in further analysis in D and E. **D)** Feature plots show extracted fibroblast subsets (cluster 4, 6, 13, 21) and the expression of selected genes. One fibroblast cluster expresses high levels of LOX and other collagen- and ECM-relevant genes. **E)** HIF-1 expression was further analyzed in fibroblasts (FB) derived from psoriasis (P, n=3) and healthy skin (H, n=3). Bar graphs show the percentage of FB that are positive (grey) and negative (while) for HIF-1.

Supplementary Figure 4 - Balsini et al.

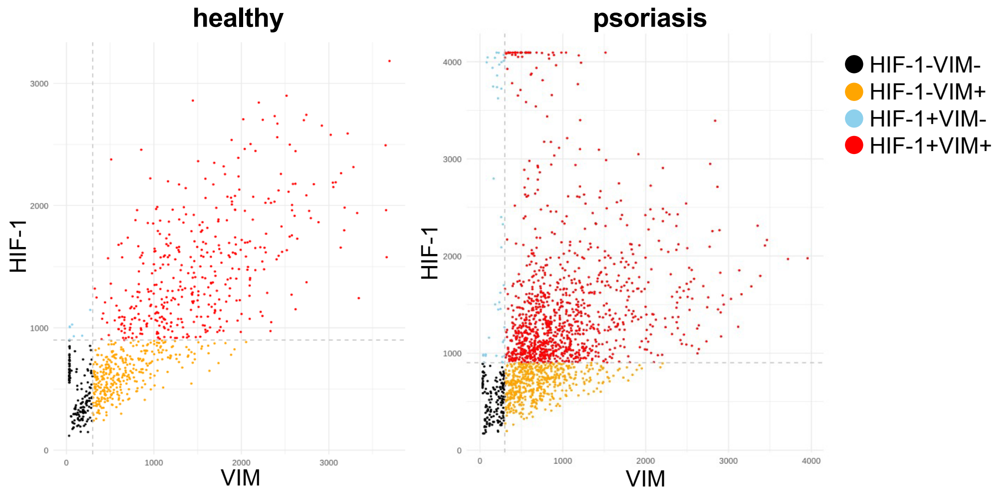


**Supplementary Figure 4. HIF-1 expression in healthy and psoriatic human skin.** Expression analysis of dermal cells extracted from immunofluorescence images of healthy and psoriatic skin. Shown are representative dot plots illustrating HIF-1 and VIM expression of extracted cells (each dot represents a cell). A gate was set to separate negative cells (black) from single positive (HIF-1+, blue; VIM+ orange) and double positive cells expressing both marker (HIF-1+VIM+, red). One representative dot plot is shown for each healthy and psoriasis. Please note the presence of a VIM+HIF-1$^{high}$ cell population in psoriasis.

Supplementary Figure 5 - Balsini et al.

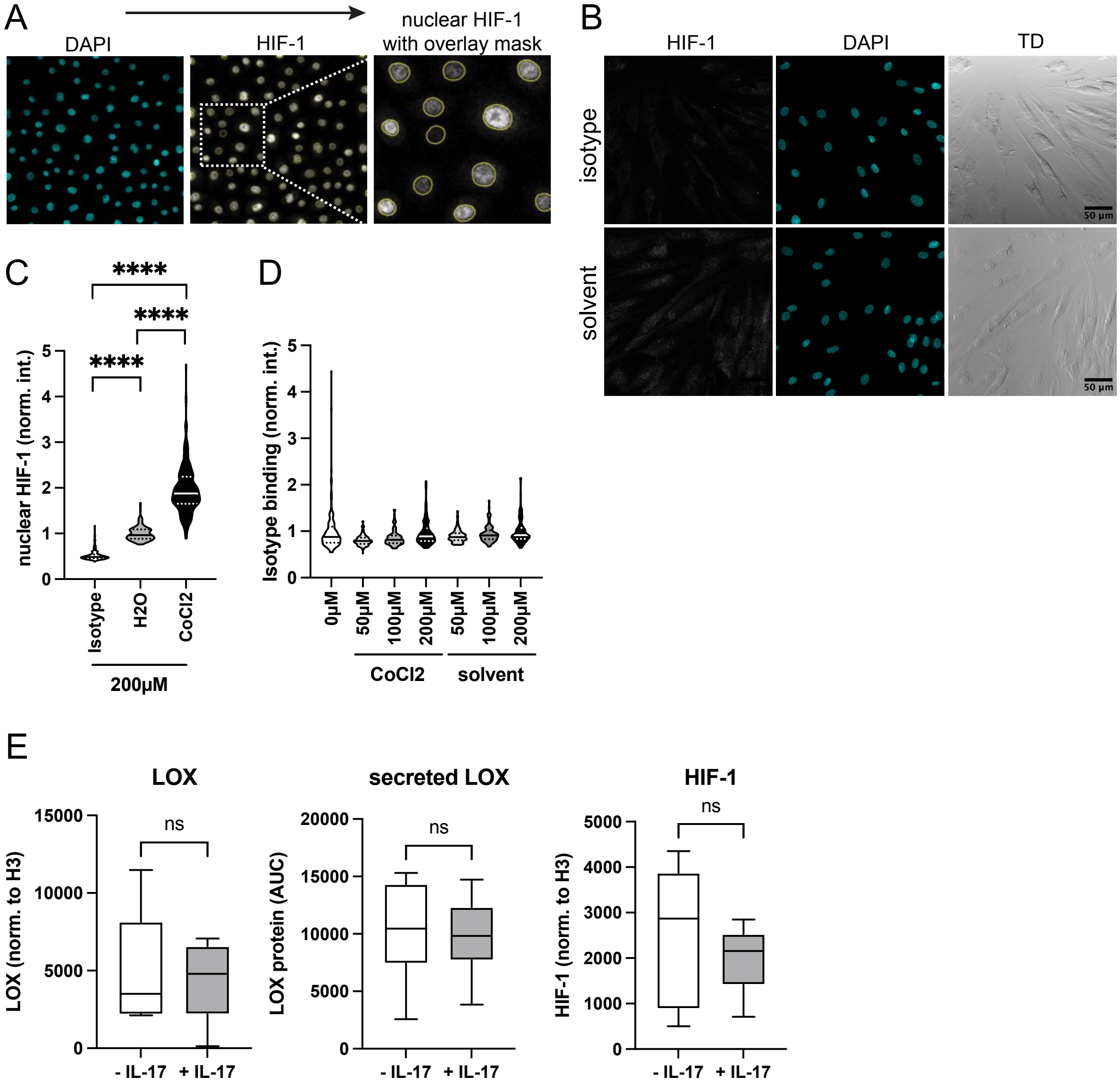


**Supplementary Figure 5. Chemical stabilization of HIF-1 with CoCl2 induces nuclear translocation and LOX secretion, while IL-17 does not. A)** Schematics illustrating steps used to analyze HIF-1 expression in nuclei. DAPI was used to mask nuclei and HIF-1 was measured in masks. **B)** Representative images of fibroblasts stained with the isotype control (upper panel) or HIF-1 upon incubation with the solvent of CoCl2 (lower panel). Scale bar = 50 µm. **C)** Quantification of nuclear HIF-1 in fibroblasts exposed to 200 µM CoCl2 or the solvent H2O. Shown is the isotype control for HIF-1, and intensity values for HIF-1 staining with and without CoCl2 incubation (n=6). **D)** CoCl2 exposure does not increase unspecific binding of antibodies. Shown are intensity value data from fibroblasts of one healthy donor

incubated with different concentrations of CoCl2 and stained with the isotype control Ab. **E)** LOX expression was evaluated upon incubation of skin fibroblast with varying concentrations of CoCl2 using qPCR (n=3). LOX mRNA was slightly, but not significantly increased. **F)** LOX expression before and upon fibroblast treatment with IL-17A. Left plot: cellular LOX.
Middle plot: secreted LOX. Right plot: HIF-1 expression before and upon fibroblast treatment with IL-17A (n=6).

Supplementary Figure 6 - Balsini et al.

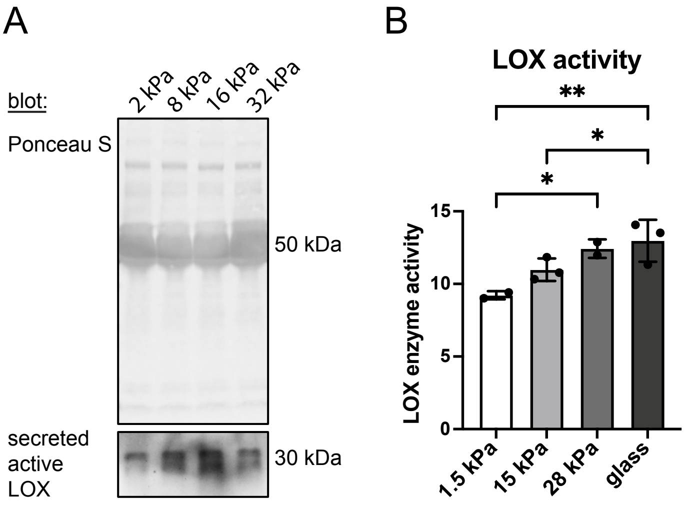


**Supplementary Figure 6. Skin fibroblasts are mechanosensitive and respond to stiff substrate with LOX secretion and enzyme activation. A)** Secreted active LOX was detected in supernatants of fibroblasts cultured on substrates with various stiffness. Ponceau S illustrates total protein content in culture supernatants. Thick bands at 50 kDa are from serum used to culture cells. Lox was detected in Western blots using antibodies against human LOX. Shown is the secreted form, which has 30 kDa. **B)** LOX activity was measured in supernatants of three fibroblast donors corresponding to stiffness values shown in Figs. 7B and 7C.
